# Decomposition of reinforcement learning deficits in gambling disorder via drift diffusion modeling and functional magnetic resonance imaging

**DOI:** 10.1101/2020.06.03.131359

**Authors:** Antonius Wiehler, Jan Peters

**Affiliations:** Department of Systems Neuroscience, University Medical Centre Hamburg-Eppendorf, Hamburg, Germany; Institut du Cerveau et de la Moelle épinière (ICM), INSERM U 1127, CNRS UMR 7225, Sorbonne Universités Paris, France; Department of Psychology, Biological Psychology, University of Cologne, Cologne, Germany

## Abstract

Gambling disorder is associated with deficits in feedback-based learning tasks, but the computational mechanisms underlying these learning impairments are still poorly understood. Here, we examined this question using a combination of computational modeling and functional resonance imaging (fMRI) in individuals that regular participate in gambling (n=23, seven fulfilled 1-3 DSM 5 criteria, sixteen fulfilled 4 or more) and matched controls (n=23). Participants performed a stationary reinforcement learning task with two pairs of stimuli (80% vs. 20% reinforcement rates per pair). As predicted, the gambling group made significantly fewer selections of the optimal stimulus, while overall response times (RTs) were not significantly different between groups. We then used comprehensive modeling using reinforcement learning drift diffusion models (RLDDMs) in combination with hierarchical Bayesian parameter estimation to shed light on the computational underpinnings of this performance impairment. In both groups, an RLDDM in which both non-decision time and response threshold (boundary separation) changed over the course of the experiment accounted for the data best. The model showed good parameter and model recovery, and posterior predictive checks revealed that in both groups, the model reproduced the evolution of both accuracy and RTs over time. The learning impairment in the gambling group was attributable to a more rapid reduction in boundary separation over time, and a reduced effect of value-differences on the drift rate, compared to controls. The gambling group also exhibted substantially shorter non-decision times. Imaging analyses replicated earlier effects of prediction error coding in the ventral striatum and value coding in the ventro-medial prefrontal cortex, but there was no credible evidence for group differences in these regions. Taken together, our findings highlight the computational mechanisms underlying reinforcement learning impairments in gambling disorder.

## Introduction

Gambling disorder is a behavioral addiction that shares core neurobiological features with substance-use-disorders [1]. Neuroimaging studies have often reported functional and to a lesser degree structural alteration in regions of the reward system in gamblers [2], in particular in the ventral striatum and the ventromedial prefrontal cortex (vmPFC), regions implicated in reinforcement learning and reward valuation [3,4]. However, as summarized extensively in a recent review [2], the directionality of dysregulation in these regions in gambling disorder is mixed, with studies reporting both increases and decreases in responses. These inconsistencies likely depend on task-specific and contextual effects [5–7], as extensively discussed previously [8–10].

Behaviorally, gambling disorder is characterized by maladaptive decision-making in a range of laboratory tasks. This includes increased temporal discounting (i.e. an increased preference for smaller-sooner over larger-later rewards) [6,11–14], and increased risk-taking [6,11,15]. There is also evidence of impairments in feedback-based learning task in disordered gambling, such as on the Wisconsin Card Sorting Test (WCST) [16–21]. Likewise, gambling disorder is associated with impairments in probabilistic reversal learning [16,22], and reductions in directed (strategic) exploration in reinforcement learning [23]. Although there is some heterogeneity across studies with respect to reversal learning impairments, the general directionality of these effects is quite consistent in the literature a [24].

Impairments in feedback-based learning might be of particular relevance in the context of gambling disorder. Modern Electronic Gaming Machines (EMGs) are designed to encourage “rapid and continuous play” [25], e.g. by combining high event frequencies with salient audio-visual reinforcers [26]. Such machine design features may then, in interaction with cognitive factors such as those outlined above (e.g. impaired reward learning, reduced self-control, increased risk-taking and perseveration) [26] give rise to a particularly rapid escalation of EMG gambling compared to more traditional forms of gambling [27].

From a computational reinforcement learning (RL) perspective [28], reversal learning impairments could arise due to changes in different processes. On the one hand, impairments could be due to response perseveration, where previous actions are repeated irrespective of learning or option values. Additionally, however, reinforcement learning requires balancing exploration (choosing options with unknown value for information gain) and exploitation (choosing options with known value for reward maximization) [28–30]. Gambling disorder is linked to a reduction in direcred exploration [23], and reversal learning impairments could therefore also be due to reduced exploration. Finally, reduced learning rates, or overall a reduced consideration of option values in the decision process could likewise underlie learning impairments in disordered gambling.

Traditionally, RL models account for trial-wise categorical decisions by assuming that choices stochastically depend on the values of the available options. These values are learned via e.g. the delta learning rule, where values are updated based on reward prediction errors [28]. This learning rule is then combined with a choice rule such as softmax action selection [28], where the slope parameter indexes the “value-dependency” or “stochasticity” of decisions with respect to the values implied by a given model. However, such choice rules are agnostic with respect to the computational processes underlying changes in “stochasticity” [31].

For this reason, recent work has begun to take the distribution of choice response times (RTs) into account. Sequential sampling models such as the drift diffusion model [32,33] (DDM) are widely used in perceptual decision-making. These models assume that choices arise from a noisy evidence accumulation process that terminates as soon the accumulated evidence exceeds a threshold. In its simplest form, the DDM has three free parameters. The drift rate *v* reflects the average rate of evidence accumulation. The boundary separation parameter α governs the response threshold, and thus controls the speed-accuracy trade-off – a smaller boundary separation emphasizes speed over accuracy, whereas the reverse is true for a larger boundary separation. The non-decision time τ models perceptual and/or motor components or the RT that are unrelated to the evidence accumulation process. If one assumes that the quality of evidence in favor of a decision option is reflected in the trial-wise drift rate (via some *linking function* [34]), the DDM can be used to model trial-wise decisions in reinforcement learning [31,35,36] and value-based decision-making more generally [37–39]. The benefits of such an approach over e.g. traditional softmax choice rules are both of technical and theoretical nature. In technical terms, inclusion of RTs during model estimation has been shown to improve parameter recovery and reliability [36,40], which is of particular relevance when the number of observations is small, e.g. when working with clinical samples. In theoretical terms, a combined reinforcement learning DDM (RLDDM) is not only a more complete model, as it accounts for both binary decisions and RTs, but also allows for a more fine-grained analysis of the dynamics underlying the decision process. For example, choices could be more random or stochastic due to a reduced impact of values on drift rates or due to a more liberal decision threshold (reduced boundary separation). RLDDM models can dissociate these different possibilities, whereas softmax choice rules only contain a single “stochasticity” parameter, and thus cannot disentangle effects of value-dependency from threshold changes. We have recently shown that DDMs can reveal alterations in decision-making following prefrontal cortex damage [37] and following pharmacological dopamine challenges [38,39,41]. Others have shown that RLDDMs can reveal pharmacological effects on learning in attention-deficit hyperactivity disorder [31] and contextual effects in reinforcement learning [35,42].

Here we extended these recently developed modeling schemes to comprehensively analyze the computational basis of reinforcement learning deficits in gambling disorder. A gambling group and a matched control group performed a simple stationary reinforcement learning task [41,43] while brain activity was measured using functional magnetic resonance imaging (fMRI). Model-agnostic analyses revealed a substantial reduction in accuracy in the gambling disorder group. Modeling using hierarchical Bayesian parameter estimation revealed that an extension of previously proposed RLDDMs [31,34,35] that allowed both non-decision times and boundary separation to vary over the course of learning accounted for the data best, showed good parameter recovery, and in both groups accounted for the evolution of accuracy and response times over the course of learning. This model revealed that reduced performance in the gambling group was linked to both a more rapid reduction in decision thresholds over time, and a reduction in the value-dependence of the drift rate. FMRI replicated core effects of prediction error and value signaling in ventral striatum and ventro-medial prefrontal cortex, but did not reveal credible group differences in these effects.

## Methods

### Participants

In total, n=23 individuals that regularly participate in gambling and n=23 matched controls participated in the study. All participants provided informed written consent prior to their participation, and the study procedure was approved by the local institutional review board (Hamburg Board of Physicians, project code PV4720).

The demographic and clinical characteristics of the groups have been reported in detail elsewhere [23]. In short, groups were matched on age (M[!SD] gamblers: 25.91 [6.47], controls: 26.58 [6.52], t=-.33, p=.74), gender (all participants were male), smoking according to the Fagerström Test for Nicotine Dependence (FTND) [44] (M[!SD] gamblers: 2.14 [2.58], controls: 2.21 [2.18], t=-.10, p=.92), alcohol use according to the Alcohol Use Disorders Test (AUDIT) [45] (M[!SD] gamblers: 6.09 [7.14], controls: 6.84 [4.91], t=-.40, p=.69) and education (school years M[!SD] gamblers: 11.64 [1.77], controls: 11.79 [1.40], t=-.31, p=.76). Gamblers scored higher on depression according to the Beck Depression Inventory (BDI-II) [46] (M[!SD] gamblers: 15.41 [11.41], controls: 8.47 [8.46], t=2.26, p=.03).

Sixteen individuals from the gambling group fulfilled four or more DSM-5 criteria for gambling disorder (previously classified as pathological gamblers). Seven individuals fulfilled one to three DSM-5 criteria (previously classified as problem gamblers). Addiction severity was further characterized using the German “Kurzfragebogen zum Glücksspielverhalten” (KFG) [47] (M[!SD] gamblers: 25.90 [14.15], controls: 0.58 [0.32], t=8.55, p<.001) and the South Oaks Gambling Screen (SOGS) [48] (M[!SD] gamblers: 8.64 [4.46], controls: 0.21 [0.54], t=8.99, p<.001). These scores were highly correlated in the gambling group (r=.95) and were thus combined for correlational analyses into a single “addiction severity” score by averaging their respective z-scores.

Due to an incidental finding during fMRI which may have impacted spatial normalization, one control participant was excluded from the imaging data analysis. This participant was however retained for all behavioral and modeling analyses. Participants were recruited via postings in local internet bulletin boards, and reported no history of neurological or psychiatric disorder except for depression. No participants were currently undergoing any psychiatric treatment. Current drug abstinence on the day of scanning was verified via a Urine drug test.

### Reinforcement learning task

Following completion of our previously reported restless four-armed bandit task [23], participants had a short break inside the scanner. Then they performed 60 trials in total of a simple reinforcement learning task [41,43] using two pairs of stimuli (n=30 trials per pair). Per pair, one stimulus was associated with a reinforcement rate of 80% (optimal stimulus) whereas the other was associated with a reinforcement rate of 20% (suboptimal stimulus). Options were randomly assigned to the left/right side of the screen, and trials from the two option pairs were presented in randomized order. Participants had three seconds to choose one of the two stimuli via button press (see Figure 1). Participants then received binary feedback, either in the form the display of a 1€ coin (*reward* feedback, see Figure 1) or as a crossed 1€ coin (*no reward* feedback). A jitter of variable duration (2-6sec, uniformly distributed) was included following presentation of the selection feedback and following presentation of the reward feedback (see Figure 1). Prior to scanning, participants performed a short practice version of the task in order to familiarize themselves with the task and the response deadline.

**Figure 1.**
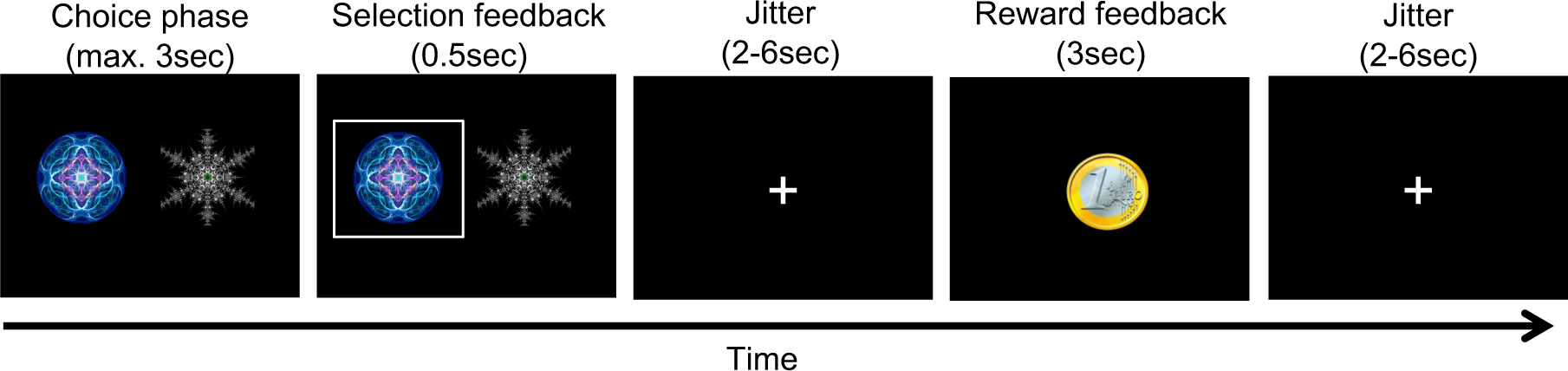
Illustration of a single trial from the reinforcement learning task. Stimuli were presented for a maximum of 3sec, during which participants were free to make their selection. The selection was then highlighted for 500ms, followed by a jitter of variable duration (2-6sec). Reward feedback was then presented for 3sec, followed by another jitter of variable duration (2-6sec). Stimuli consisted of two pairs of abstract fractal images (80% vs. 20% reinforcement rate), which were presented in randomized order, and participants completed 30 trials per pair.

### Model-agnostic statistical analyses

Model-agnostic measures (accuracy, median RT) where analyzed using Bayesian Wilcoxon Rank-Sum tests as impletemented in JASP [49] (Version 0.l6.3).

### Q-learning model

We applied a simple Q-learning model [28] to formally model the learning process. Here, participants are assumed to update the value (Q-value, Eq. 1) of the chosen action *i* based on the reward prediction error that is computed on each trial as the difference between the obtained reward and the expected reward (Eq. 2):

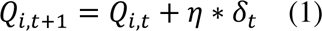

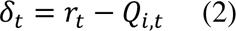

Q-values of unchosen actions remain unchanged. All Q-values were initialized with values of 0.5. As learning from positive and negative feedback is thought to depend on distinct striatal circuits [50,51], also examined models with separate learning rates (η_+_, η_-_) for positive vs. negative prediction errors. Learning rates were estimated in standard normal space [-3, 3] and back-transformed to the interval [0, 1] via the inverse cumulative normal distribution function.

### Softmax action selection

Softmax action selection models the choice probability of the chosen action *i* on trial *t* as a sigmoid function of the Q-value difference [52] between the optimal and suboptimal options:

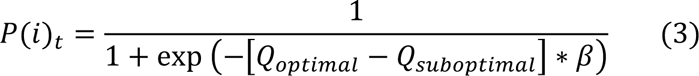

The inverse temperature parameter β models the degree to which choice probabilities depend on Q-values, such that choices are random for β = 0, and increasingly depend on the Q-value differences between options as β increases.

### Reinforcement learning drift diffusion models (RLDDMs)

We next set out to more comprehensively analyze choice dynamics underlying learning performance. To this end, we examined a set a reinforcement learning drift diffusion models [31,35,41] (RLDDMs) in which the DDM replaces softmax action selection as the choice rule [34]. These models can account for the full response time (RT) distributions associated with decisions, and thus provide additional information regarding the dynamics of the choice process.

The upper response boundary was defined as selection of the optimal (80% reinforced) stimulus, whereas the lower response boundary was defined as selection of the suboptimal (20% reinforced) stimulus. RTs for choices of the suboptimal option where multiplied by -1 prior to model estimation, and we discarded for each participant the fastest 5% of trials. In a null model without a learning component (DDM_0_), the RT on each trial *t* is then distributed according to the Wiener First Passage Time (*wfpt*):

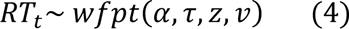

Here the boundary separation parameter α regulates the speed-accuracy trade-off, such that smaller values of α lead to faster but less accurate responses. The drift rate *v* reflects the quality of the evidence, such that greater values of *v* give rise to more accurate and faster responses. Note that in this model *v* is constant and not affected by the learning process. The non-decision time τ models RT components related to motor and/or perceptual processing that are unrelated to the evidence accumulation process. The starting point parameter *z* models a bias towards one of the response boundaries. We fixed *z* at .5 as options were presented in randomized order on the left vs. right side of the screen, and an *a priori* bias towas optimal or suboptimal choices is not plausbile in this learning setting.

Following earlier work [31] we then incorporated the learning process (Equations 1 and 2) in the DDM by setting trial-wise drift rates to be proportional to the difference in Q-values between optimal and suboptimal options using a simple linear linkage function [31,34,41]:

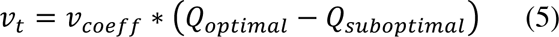

*v*_.&/00_ models the degree to which trial-wise drift rates scale with the value difference between options. The intuition is that as Q-value differences increase, the accuracy increases and RTs decrease. Conversely, when cue-values are similar (and thus response conflict is high) choices are both noisier and slower. Note that we also examined a non-linear mapping scheme proposed earlier [35], but, as in related work [41] these models failed to converge in our data. This is likely attributable to the lower trial numbers in the present study compared to previous implementations of non-linear drift rate scaling [35,37,38].

We also examined two further extensions of the RLDDM that might capture additional RT effects unrelated to the learning process. These extensions were motiveated by the observation that in the gambling group, RTs decreased over the course of the experiment, but this effect appeared to be only in part attributable to learning, such that models with constant α and τ did not reproduce RT changes in the gambling group (see posterior predictive checks below). Therefore, we examined whether allowing boundary separation [31,35] and/or non-decision-time to vary over the course of the experiment according to a power function could account for these effects, such that

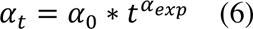

for the case of boundary separation that varies across trials *t*, and

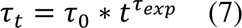

for the case of non-decision times that vary across trials *t*. On trial 1, boundary separation and non-decision time correspond to α_1_ and τ_1_. Values then change over trials according power functions with exponents α_/4’_ and τ_/4’_. The first case (Eq. 6) captures the idea that, over time, participants response thresholds might decrease due to e.g. impatience, fatigue or boredom with the task. The second case (Eq. 7) corresponds to the idea that motor and/or perceptual processes might speed up over time, e.g. due to practice effects or increased familiarity with the task.

Therefore, the model space included the null model (DDM_0_) and eight variants of the RLDDM, which differed according to learning rates (single vs. dual), decision thresholds (fixed vs. power function) and non-decision times (fixed vs. power function).

### Hierarchical Bayesian models

Models were fit to all trials from all participants, separate for each group, using a hierarchical Bayesian modeling approach with group-level Gaussian distributions for all parameters. Posterior distributions were estimated using Markov Chain Monte Carlo as implemented in the JAGS software package [53] (Version 4.3) using the Wiener module for JAGS [54] distribution, in combination with Matlab (The MathWorks) and the *matjags* interface (https://github.com/msteyvers/matjags). For group-level means and standard deviations, we defined uniform priors over numerically plausible parameter ranges (see Table 1).

**Table 1.** Overview of priors for group means.

| Parameter | Group-level prior ($\mu$) | Group-level prior ($\sigma$) |
| --- | --- | --- |
| $\alpha_0$ | *Uniform* (.01, 5) | *Uniform* (.0001, 2) |
| $\alpha_{\text{exp}}$ | *Uniform* (-3, 3) | |
| $\tau_0$ | *Uniform* (0.1, 2) | *Uniform* (.0001, 2) |
| $\tau_{\text{exp}}$ | *Uniform* (-3, 3) | |
| $v_{\text{coeff}}$ | *Uniform* (-100, 100) | *Uniform* (.0001, 10) |
| $\eta_+, \eta_-$ | *Uniform* (-3, 3) | *Uniform* (.0001, 4) |

For each model and group, we ran 2 chains with a burn-in period of 50.000 samples and thinning factor of 2. 10.000 additional samples were then retained for further analysis. Chain convergence was assessed by examining the Gelman-Rubinstein convergence diagnostic 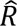, and values of 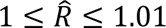 were considered as acceptable for all group-level and individual-subject parameters. Relative model comparison was performed via the Widely Applicable Information Criterion (WAIC) and the estimated log pointwise predictive density (*elpd*) [55], an approximation of the leave-one-out cross-validation accuracy of the model.

### Parameter recovery simulations

Parameter recovery simulations were conducted to ensure that known parameters underlying the data-generating process could be recovered using our modeling procedures. For this purpose, we simulated 10.000 full data sets from the posterior distribution of the best-fitting models, randomly selected 10 of these simulated data sets, and re-fit the models to these simulated data. Parameter recovery was then assessed via two procedures. For subject-level parameters, we examined the correlation between generating and estimated parameters across all ten simulations. For group-level means and standard deviations, we examined whether the estimated 95% highest posterior density intervals contained the true generating parameter value.

### Model recovery simulations

To ensure that the true data-generating model could be identified using our modeling procedures, model recovery analyses were conducted, focusing on the three best-fitting models (RLDDMs 4, 6 and 8). N=20 full datasets were simulated from each of the three models’ posterior distributions, and re-fit with all nine models from the model space. The percentage of simulations in which the true data-generating model was recovered was then taken as a measure of model recovery.

### Posterior predictive checks

Posterior predictive checks were performed to ensure that the best-fitting models captured key aspects of the data, again using data sets simulated from each model’s posterior distributions. For each simulated data set, we then computed for each group the mean RTs and accuracies for 10 trial bins (averaging across 1000 randomly selected simulated data sets), and compared these model-predicted values to the observed data per group. Individual-participant posterior pedictive checks were carried out by overlaying simulated and observed individual-participant RT distributions, and by overlaying simulated and observed RT changes over the course of learning via five trial bins.

### Analyses of posterior distributions

Posterior distributions are reported in the following ways. Mean group differences along with 95% highest density intervals are examined, and the posterior probabilities for group differences > 0 are reported. For completeness, we also report directed Bayes Factors that reflect the relative evidence in favour of a group difference < 0 compared vs. a group difference > 0.

### FMRI data acquisition

MRI data were collected on a Siemens Trio 3T system using a 32-channel head coil. Participants performed a single run of 60 trials in total (following a short break, after completion of our previously reported task [23]). Each volume consisted of 40 slices (2 x 2 x 2 mm in-plane resolution and 1-mm gap, repetition time = 2.47s, echo time 26ms). We tilted volumes by 30° from the anterior and posterior commissures connection line to reduce signal drop out in the ventromedial prefrontal cortex and medial orbitofrontal cortex [56]. Participants viewed the screen via a head-coil mounted mirror, and logged their responses via the index and middle finger of their dominant hand using an MRI compatible button box. High-resolution T1 weighted structural images were obtained following completion of the cognitive tasks.

### FMRI preprocessing

All preprocessing and statistical analyses of the imaging data was performed using SPM12 (Wellcome Department of Cognitive Neurology, London, United Kingdom). As in our previous study in this sample [23], volumes were first realigned and unwarped to account for head movement and distortion during scanning. Second, slice time correction to the onset of the middle slice was performed to account for the shifted acquisition time of slices within a volume. Third, structural images were co-registered to the functional images. Finally, all images were smoothed (8mm FWHM) and normalized to MNI-space using the DARTEL tools included in SPM12 and the VBM8 template.

### FMRI statistical analysis

Error trials were defined as trials were no response was made, or trials that were excluded from the computational modeling during RT-based trial filtering (see above, recall that for each participant, the fastest 5% of trials were excluded).

Following earlier work [41], three first-level general linear models (GLMs) were examined. GLM1 used the following regressors:

1. onset of the decision option presentation
2. onset of the decision option presentation modulated by chosen – unchosen Q-value
3. onset of the decision option presentation modulated by (chosen – unchosen Q-value)^2^
4. onset of the feedback presentation
5. onset of the feedback presentation modulated by model-based prediction error
6. onset of the decision option presentation for error trials
7. onset of the feedback presentation for error trials.

In GLM2, chosen – unchosen value was replaced with the average Q-value across options. GLM3 used the following regressors:

1. onset of the decision option presentation
2. onset of the decision option presentation modulated by chosen – unchosen value
3. onset of the decision option presentation modulated by (chosen – unchosen value)^2^
4. onset of the feedback presentation for positive prediction errors
5. onset of the feedback presentation for negative prediction errors
6. onset of the feedback presentation for error trials.

Following earlier work using this task [41,43], Q-values and prediction errors were computed using the posterior group-mean learning rates from the best-fitting final hierarchical Bayesian model (RLDDM8). Parametric modulators were *z*-scored within-subjects prior to entering them into the first level model [57]. Single-subject contrast estimates were then taken to a second-level random effects analysis using the two-sample t-test model as implemented in SPM12. At the second level, the following *z*-scored covariates were included: age, depression as assessed via the Beck Depression Inventory II (BDI) [46], smoking behavior as assessed via the Fagerström Test for Nicotine Dependence (FTND) [44] and alcohol use as assessed via the Alcohol Use Disorders Identification Test (AUDIT) [45].

All contrasts are displayed at p<.001 (*uncorrected*) with k>=10 voxels, and correction for multiple comparisons using the family-wise error rate (FWE) followed the same approach as in our earlier work [41] and used a sigle region-of-interest (ROI) mask provided by the Rangel Lab (https://www.rnl.caltech.edu/resources/index.html) that is based on two meta-analysis of reward valuation effects [3,4]. This mask covers core areas involved in reward processing, including bilateral ventral striatum, ventromedial prefrontal cortex, anterior cingulate cortex and posterior cingulate.

## Results

Behavioral data analysis and computational modeling proceeded in the following steps. We first analyzed model-free performance measures. Next, we carried out a detailed model comparison of a set of candidate reinforcement learning drift diffusion models (RLDDMs) and identified the best-fitting models per group. We then ran parameter recovery analyses to ascertain that the true data-generating parameters could be recovered, and ran posterior predictive checks to ensure that the key patterns in the data could be reproduced by the best-fitting models. Finally, we examined the parameter posterior distributions and compared them between groups, before moving to the analysis of the fMRI data.

**Figure 1.**
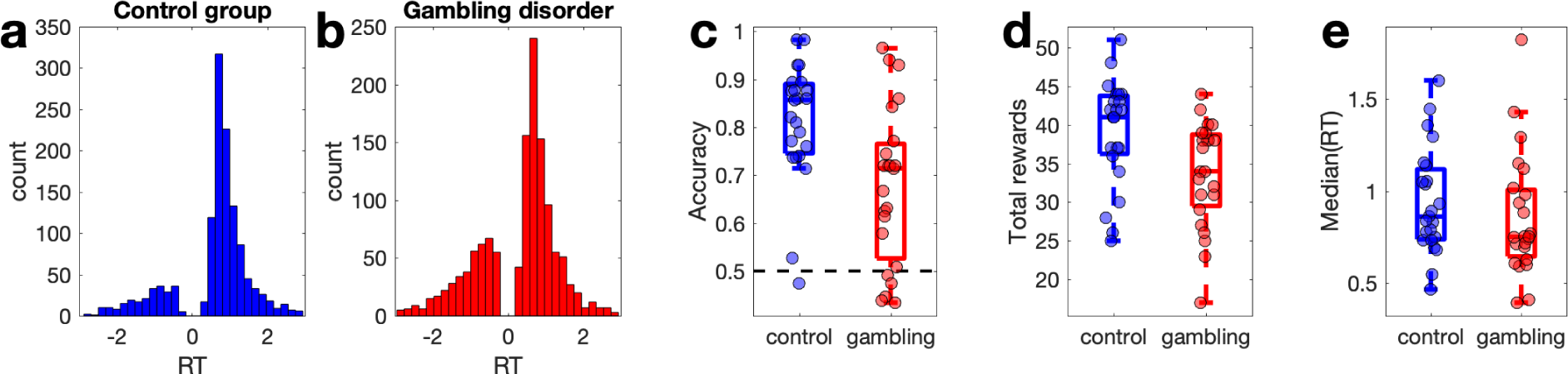
Response time (RT) distributions the control group (a, blue) and the gambling disorder group (b, red) with choices of the suboptimal options coded as negative RTs. c: Accuracy per group (chance level is 0.5). d: Total rewards earned per group. e: Median RTs per group.

### Model-agnostic analysis

RT distributions per group are shown in Figure 1a and 1b, with choices of the suboptimal option coded as negative RTs. While control group participants selected the optimal stimulus on around 80% of trials (Figure 1c), gambling group participants only made around 68% correct choices. A Bayesian Wilcoxon Rank sum test confirmed moderate evidence for a group difference in accuracy (BF10 = 6.67, Figure 1c) and total reward obtained (BF10 = 3.94, Figure 1d). For median RTs, in contrast, a Bayesian Wilcoxon Rank Sum test revealed anecdotal evidence for the null model (BF01 = 1.87, Figure 1e).

### Model comparison

We next compared a range of computational models (see methods section). As a reference, we first fit a null model (DDM_0_) without a learning component. Next, a set of reinforcement learning DDMs (RLDDMs) was examined that all included a linear mapping from Q-value differences to trial-wise drift rates [31,34,41] (see Eq. 5). This modeling scheme incorporates the intuition that successful learning should decrease RTs and increase accuracies, and that accuracy should be higher and RTs shorter when making easier choice (i.e. when Q-value differences are larger). The model space included models with single vs. dual learning rates η (for positive vs. negative prediction errors), and models with fixed vs. modulated boundary separation α and non-decision times τ (see Eq. 6 and 7), yielding a total of eight RLDDMs (see Table 2).

**Table 2.** Model comparison results, separately per group. We examined reinforcement learning drift diffusion models (RLDDMs) with single vs. dual learning rates (η) and fixed vs. modulated non-decision times (τ) and boundary separation (α), as well as a null model without learning (DDM_0_). Model comparison used the estimated log pointwise predictive density (-elpd)[55]. We also report the 95% CI of the difference in -elpd between each model and the best-fitting model (-elpd_diff_).

| Model | $\eta$ | $\tau$ | $\alpha$ | Controls | | | Gamblers | | |
| --- | --- | --- | --- | --- | --- | --- | --- | --- | --- |
|  |  |  |  | -elpd | -elpd<sub>diff</sub> | Rank | -elpd | -elpd<sub>diff</sub> | Rank |
| DDM<sub>0</sub> | - | Fixed | Fixed | 800.2 | 215.0<br>[171.6, 258.4] | 9 | 1115.9 | 107.6<br>[78.9, 136.3] | 9 |
| RLDDM1 | 1 | Fixed | Fixed | 658.1 | 72.8<br>[48.6, 97.1] | 8 | 1055.5 | 47.3<br>[27.3, 67.2] | 8 |
| RLDDM2 | 1 | Fixed | Power | 634.3 | 49.0<br>[29.3, 80.7] | 6 | 1021.4 | 13.1<br>[.4, 25.8] | 4 |
| RLDDM3 | 1 | Power | Fixed | 644.0 | 58.8<br>[36.8, 97.1] | 7 | 1027.1 | 18.8<br>[3.1, 34.4] | 6 |
| RLDDM4 | 1 | Power | Power | 628.3 | 43.1<br>[24.1, 68.7] | 5 | 1010.3 | 2.0<br>[-9.1, 13.2] | 2 |
| RLDDM5 | 2 | Fixed | Fixed | 615.0 | 29.7<br>[15.9, 43.5] | 4 | 1049.0 | 40.7<br>[23.8, 57.6] | 7 |
| RLDDM6 | 2 | Fixed | Power | 591.4 | 6.1<br>[-.1, 12.3] | 2 | 1019.3 | 11.1<br>[4.5, 17.6] | 3 |
| RLDDM7 | 2 | Power | Fixed | 599.8 | 14.5<br>[4.4, 24.6] | 3 | 1022.4 | 14.1<br>[3.4, 24.9] | 5 |
| RLDDM8 | 2 | Power | Power | 585.3 | 0.0 | 1 | 1008.3 | 0.0 | 1 |

Model comparison was performed using the estimated log pointwise predictive density (-elpd) [55] (Table 2). In both groups, the data were generally better accounted for by models in which both α and τ were modulated according to a power function, and in both groups, RLDDM8 exhibited lowest -elpd value. However, the 95% confidence intervals of the -elpd difference between the best model and the second-best model (RLDDM6 in the control group and RLDDM4 in the gambling group) overlapped with zero, indicating that the evidence in favour of RLDDM8 was overall not decisive.

Despite this inconclusive model comparison, we focused all remaining analyses on RLDDM8, for the following reasons. First, in the control group, the overlap in -elpd between RLDDM6 and 8 was numerically very small. Second, model recovery was substantially better for RLDDM8 than RLDDM4 and RLDDM6 (see below). Third, RLDDM4 and 6 are nested versions of RLDDM8. In RLDDM4, positive and negative learning rates are identical, η_+_ = η_-_, and in RLDDM6, τ_/4’_= 0. In such cases an estimation approach (i.e. examining the posterior distributions of the parameters) may be more informative than relying solely on categorical model comparison [58]. The reason is that a parameter’s posterior distribution provides the best information regarding the value of a parameter, given the priors and the data, and thus allows for a quantification of the degree of evidence that e.g. learning rates differ, or that τ_/4’_ is different from 0.

### Parameter and model recovery simulations

Parameter recovery analyses were carried out across 10 simulated datasets. Results are provided in Supplemental Figure 1 for RLDDM8 and Supplemental Figure 2 for RLDDM4. All correlations between generating and estimated individual-subject parameters were ≥.57 (see Supplemental Table 1) and group-level parameters recovered well (Supplemental Figures 1 and 2).

Model recovery analyses were restricted to the best fitting model (RLDDM8) and the two runner-up models (RLDDM4 and RLDDM6). Amonst these models, RLDDM8 exhibited the best model recovery performance (see Supplemental Figure 3), such that in 77% of simulations from RLDDM8, this model also provided the best fit amongst all models from the model space.

### Posterior predictive checks

As a model comparison is always relative to a given set of candidate models, we next performed posterior predictive checks to examine the degree to which RLDDM8 accounted for key patterns in the data, in particular with respect to changes in accuracy and RT over the course of learning. For comparison, we included the DDM_0_, and the simplest learning model (RLDDM1), and overlayed mean accuracies and RTs per time bin of simulated data and observed data (see methods section), separately for each group. Figure 3 (control group) and Figure 4 (gambling group) depict the observed and model-predicted accuracies and RTs per trial bin.

**Figure 3.**
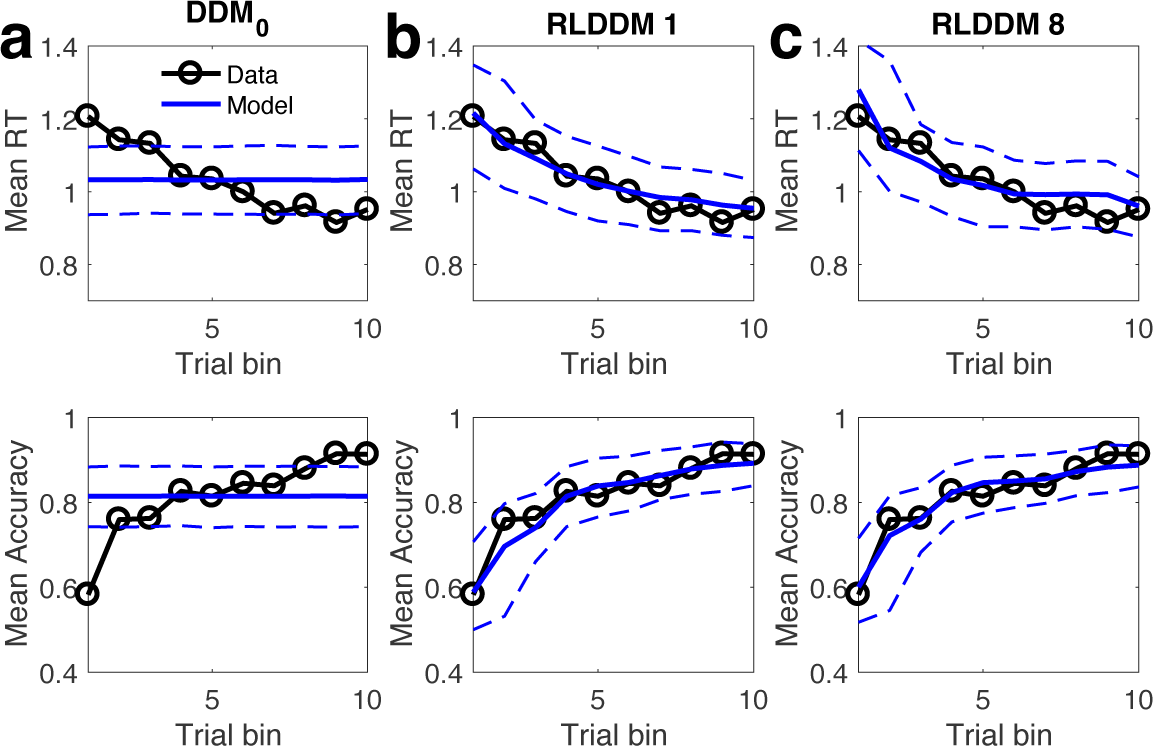
Posterior predictive checks in the control group. Top row: observed RTs over time (black lines) and model predicted RTs (solid blue lines: means, dashed lines: +/-95% percentiles). Bottom row shows observed accuracies over time (black lines) and model predicted accuracies (solid blue lines: means, dashed lines: +/-95% percentiles). a) DDM0 without reinforcement learning. b) RLDDM1 with a single learning rate, fixed non-decision time and fixed boundary separation. c) RLDDM8 with dual learning rates, modulated non-decision time and modulated boundary separation.

**Figure 4.**
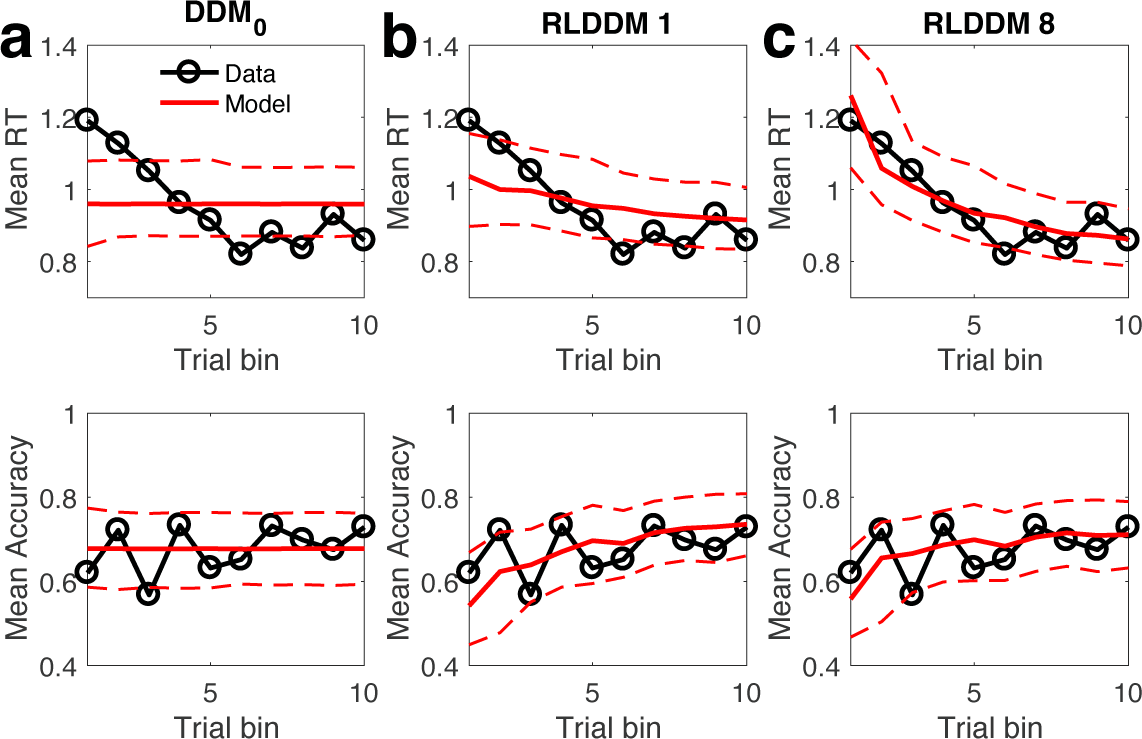
Posterior predictive checks in the gambling disorder group. Top row: observed RTs over time (black lines) and model predicted RTs (solid red lines: means, dashed lines: +/-95% percentiles). Bottom row shows observed accuracies over time (black lines) and model predicted accuracies (solid red lines: means, dashed lines: +/-95% percentiles). a) DDM0 without reinforcement learning. b) RLDDM1 with a single learning rate, fixed non-decision time and fixed boundary separation. c) RLDDM8 with dual learning rates, modulated non-decision time and modulated boundary separation.

DDM_0_ predicts constant accuracies and RTs over trials, and as can be seen in Figures 3a and 4a, it cannot reproduce predicted learning-related changes. In contrast, RLDDMs predict learning-related increases in accuracy and decreases in RTs over time. Notably, in the control group (Figure 3b,c), both RLDDM1 and RLDDM8 provide a reasonbly good account of both effects on the group level. In contrast, in the gambling disorder group (Figure 4b, c), RLDDM1 provided a poor account of group-level changes in RTs over time, suggesting that reinforcement learning alone was insufficient to account for the RT reductions over time in the gambling group.

Individual-participant posterior predictive checks confirmed that RLDDM8 provided a good account of individual-participant RT distributions (Supplemental Figures 4 and 5) and the change in RTs over the course of learning in individual subjects (Supplemental Figures 6 and 7).

### Group differences in model parameters

Next, group differences differences in RLDDM8 parameters were examined in detail. Posterior distributions of parameter group means as well as group differences are shown in Figure 5 for each RLDDM parameter. Analyses of posterior distributions are provided in detail in Table 3. Three reliable group differences emerged, with posterior probabilities >96% (Table 3): First, α_exp_ was reliably reduced in the gambling group compared to the control group (Figure 3b and Table 3, model-implied boundary separation changes over time for each group are shown in Supplemental Figure 8). The gambling disorder group thus showed a more rapid reduction in decision thresholds over time than the control group. Second, the offset of the non-decision-time, τ_0_, was reliably lower in the gambling group compared to the control group (Figure 3c and Table 3). Third, the drift rate value modulation, *v_coeff_*, was reliably lower in the gamling group compared to the control group (Figure 3e and Table 3).

**Figure 5.**
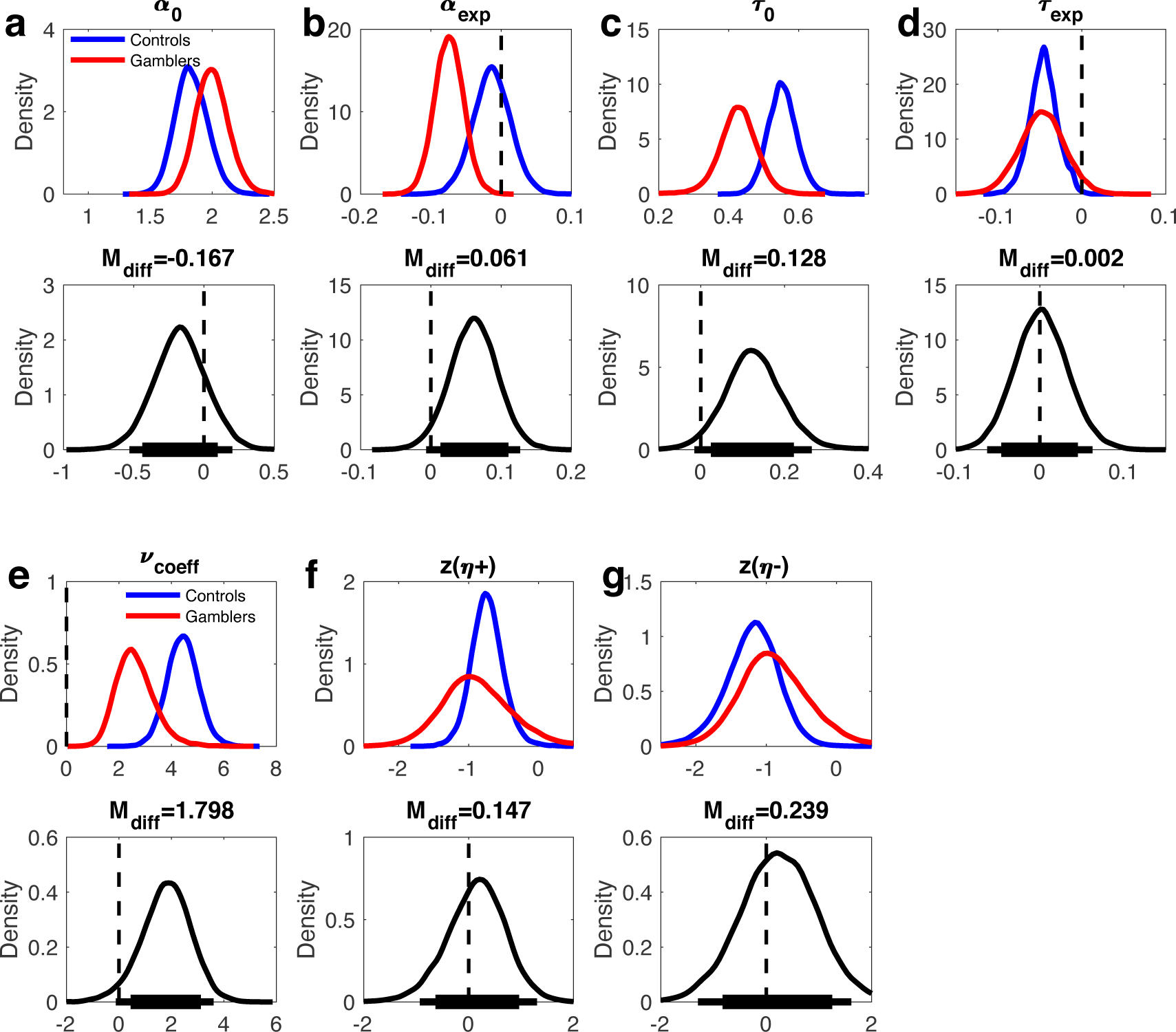
Posterior distributions for RLDDM8 parameters. Upper panels: posterior distributions of parameter group means for the control group (blue) and the gambling group (red). Lower panels: posterior group differences per parameter (controls group – gambling disorder group). Solid (thin) horizontal lines in the lower panels denote 85% (95%) highest posterior density intervals.

**Table 3.** Group differences and within-group effects for all RLDDM8 parameters. M_diff_: mean posterior group difference. P(group diff. > 0): posterior probability that the group difference in a parameter is > 0. dBF (group difference): directional Bayes Factors comparing the evidence for a group difference > 0 to the evidence for a group difference < 0. Within group comparisons: P(effect): posterior probability that for an effect (for α_exp_ and τ_exp_, the comparison is against 1, for *v_coeff_* the comparison is against 0). dBF: directional Bayes Factors comparing the evidence for a parameter value > 0 to the evidence for a parameter value < 0.

|  | Group differences |  |  | Within-group comparisons |  |  |  |
| --- | --- | --- | --- | --- | --- | --- | --- |
| | $M_{diff}$ | $P(\text{group diff.} > 0)$ | $dBF$ | Control group | | Gambling group | |
| | | | | $P(\text{effect})$ | $dBF$ | $P(\text{effect})$ | $dBF$ |
| $\alpha_0$ | -.167 | 18.25% | .29 | | | - | |
| $\alpha_{exp}$ | .061 | 96.39% | 27.60 | 69.29% | .43 | 99.98% | .00024 |
| $\tau_0$ | .128 | 96.91% | 27.09 | | | - | |
| $\tau_{exp}$ | .002 | 52.05% | 1.12 | 99.71% | .003 | 96.10% | .045 |
| $v_{coeff}$ | 1.79 | 96.40% | 25.79 | >99.99% | 15860 | >99.99% | 15828 |
| $\eta_+$ | .147 | 62.44% | 1.64 | | | - | |
| $\eta_-$ | .239 | 63.09% | 1.66 | | | - | |

For comparison, behavioral data were also fitted with a standard softmax choice rule (Equation 3). Here, the inverse temperature parameter (β) was substantially reduced in the gambling group compared to the control group (see Supplemental Figure 9 and Supplemental Table 2). This result is consistent with the effects observed for the RLDDM8, as both a lower value coefficient of the drift rate and a lower boundary separation would translate to higher levels of decision noise in the softmax model.

### FMRI results

In a first step, replication analyses for previously reported effects were conducted, focusing on model-based chosen - unchosen value and model-based prediction error (based on GLM1 and GLM3) and model-based average Q-value (GLM2). We focused on a single ROI covering areas linked to reward valuation effects based on two meta-analyses (see methods section). This revealed significant main effects across groups for average Q-values in the ventromedial prefrontal cortex (Figure 6 and Table 4). A Bayesian two-sample t-test revealed moderate evidence in favour of the absence of a group effect (BF_01_=3.179).

**Figure 6.**
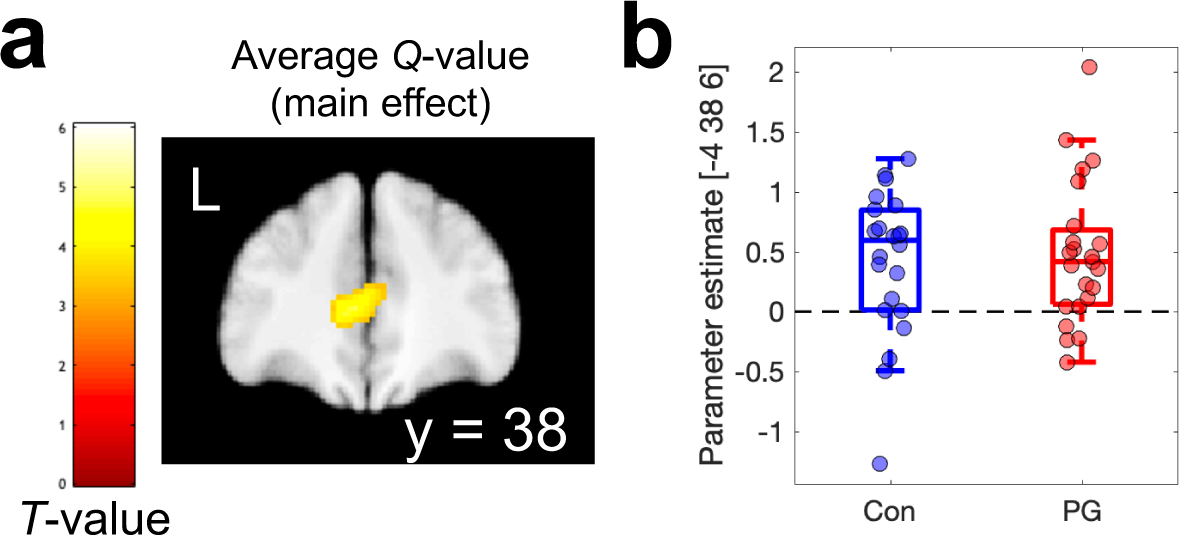
Parametric analyses of model-based average Q-values (GLM2) revealed a robust main effect across groups in the ventro-medial prefrontal cortex (a). Parameter estimates at the peak voxel from (a) are shown in b).

**Table 4.** Replication analyses for model-derived measures (main effects across groups): average Q-value across options, chosen – unchosen Q-value, and model-derived prediction error. Small volume correction for multiple comparisons (SVC) used a single region of interest mask across two meta-analyses[3,4] of reward value effects (see methods section).

| Contrast / Region | Coordinates | Peak T-value | $p(\text{FWE})_{\text{svc}}$ |
| --- | --- | --- | --- |
| Average Q-value |  |  |  |
| *vmPFC* | -4 38 6 | 4.73 | .002 |
| Chosen-unchosen value |  |  |  |
| *No significant effects in ROI* |  |  |  |
| Reward prediction error |  |  |  |
| *Left ventral striatum* | -10 6 -10 | 5.69 | <.001 |
| *Right ventral striatum* | 12 10 -12 | 6.77 | <.001 |
| *vmPFC* | -4 56 -4 | 6.26 | <.001 |
| *Posterior Cingulate Cortex* | 0 -36 -36 | 4.49 | .012 |

There were significant effects of model-based prediction errors in bilateral ventral striatum, ventromedial prefrontal cortex and posterior cingulate cortex (Figure 7 and Table 5), whereas no significant effects were observed for chosen – unchosen Q-values in our ROI. Prediction error effects were first identified via a parametric modulator in GLM1, and then visualized by extracting separate parameter estimates for positive vs. negative prediction errors from GLM3 (Figure 7b). Statistical analysis of group differences at peak voxels showing main effects Bayesian repeated measures ANOVAs with the within-subjects factor prediction error (positive/negative) and the between-subjects factor group (gambling/control). In both left and right ventral striatum, this revealed decisive evidence for an effect of prediction error, but only inconclusive evidence for overall group effects (BF_incl_ < 1) and only anecdotal evidence for the presence of group x prediction error interactions (1 < BF_incl_ < 3, see Table 5).

**Figure 7.**
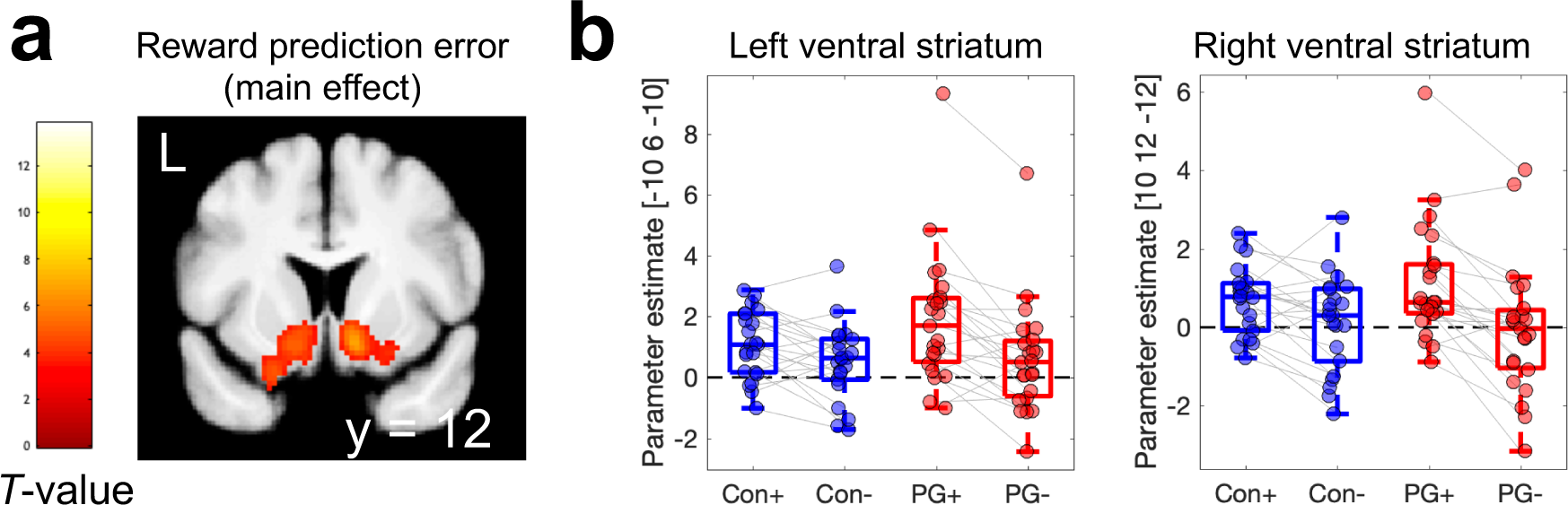
Parametric analysis of model-based reward prediction error (GLM1) revealed a robust main effect across groups in bilateral ventral striatum (a). Parameter estimates at peak voxels in (a) were then extracted from GLM3 to illustrate effects of positive (+) vs. negative (-) prediction errors in each group in both left and right ventral striatum (b).

**Table 5.** Inclusion Bayes Factors (BF_incl_) from Bayesian repeated measures ANOVAs at ventral striatal peak voxels showing main effects of model-based prediction error (PE) across groups (see Table 4).

| Effects | Left ventral striatum<br>[-10 6 -10] | Right ventral striatum<br>[10 12 -12] |
| --- | --- | --- |
| PE sign | 1060.335 | 7059.337 |
| Group | 0.878 | 0.719 |
| Group * PE Sign | 1.966 | 1.759 |

## Discussion

Here we comprehensively examined the computational underpinnings of reinforcement learning impairments in a gambling group (n=23, n=7 fulfilling one to three DSM 5 criteria for gambling disorder, and n=16 fulfilling four or more criteria) and a matched control group (n=23), using a combination of computational modeling and functional magnetic resonance imaging (fMRI). Accuracy on the learning task was substantially reduced in the gambling group, whereas there was little credible evidence for group differences in response times (RTs). Computational modeling revealed that in both groups, extended reinforcement learning drift diffusion models (RLDDMs) in which both non-decision time and boundary separation (decision threshold) were modulated according to a power function provided a superior account of the data (see below for discussion). The model with the best numerical fit (RLDDM8) showed good parameter and model recovery, and accurately reproduced the observed accuracy and response time (RTs) changes over the course of learning. Computational modeling then revealed three major group differences. Compared to the control group, the gambling group exhibited shorter non-decision times, a more rapid reduction of boundary separation (decision thresholds) over the course of learning, and a reduced value modulation of the drift rate. Neuroimaging analyses replicated effects of value in ventromedial prefrontal cortex, and prediction error in ventral striatum. However, Bayesian analyses showed that evidence for group differences in these effects was at most anecdotal.

Model comparison showed that, numerically, RLDDM8 (a model with dual learning rates, and modulated non-decision time and boundary separation according to power functions) exhibited the best fit in both groups. However, there was some overlap in the 95% confidence interval of the -elpd difference between RLDDM8 and second best model in both groups. The runner-up model additionally differed between groups (RLDDM6 for the control group, and RLDDM4 for the gambling group), such that, overall, the model comparison somewhat inconclusive. We nonetheless decided to focus subsequent analyses on the RLDDM8, with the following reasoning. Most importantly, the runner-up models in both groups are nested versions of RLDDM8. In such cases, an estimation approach (rather than categorical model comparison) can be more informative [58], as it allows a quantification of the degree of evidence that e.g. learning rates differ, or that the non-decision time modulation (τ_exp_) is different from 0. Indeed, despite the inconclusive model comparison, τ_exp_ was reliably < 0 in both groups..

We performed extensive checks to verify the performance of the non-decision-time and boundary modulation extentions in RLDDM8. First, we ran a series of parameter recovery simulations, which revealed that both subject-level and group-level parameters recovered well. Parameter recovery essentially determines the upper bound of reliability. It is therefore reassuring that estimated subject-level parameters showed a correlation between .59 and .90 with the true generating parameters. Likewise, estimated posterior distributions of group-level parameters generally contained the true generating parameters within their 95% highest posterior density intervals [35]. RLDDM8 also showed satisfactory model recovery performance, which was numerically better than both RLDDM4 and RLDDM6. Second, model performance was verified in a series of posterior predictive checks. In both groups, RLDDM8 reproduced both the increases in accuracy and the decreases in RTs over trials well. The requirement of including modulated decision thresholds was particularly evident in the gambling group, where simpler models without modulated boundary separation failed to fully account for the reductions in RTs over trials. RLDDM8 also reproduced both individual-participant RT distributions as well as RT changes over trials in individual participants.

Analysis of model parameters then allowed us to examine group differences in computational processes underlying task performance. Non-decision times, reflecting aspects of the RT that are unrelated to the evidence accumulation process, showed a similar decay over time in both groups, but the non-decision time offset τ_0_ was substantially lower in the gambling group. In contrast, the boundary separation showed a substantially more rapid decay in the gambling vs. the control group (α_exp_ was reliably more negative). That is, over the course of the experiment, individuals in the gambling group, more than controls, increasingly shifted their focus from accuracy to speed. These findings converge with previous observations of increased motor impulsivity [59], higher urgency/reduced premeditation [60] and higher levels of temporal discounting in gambling disorder [11]. In addition to these alterations in the speed-accuracy trade-off, performance deficits in the gambling group were linked to a substantial reduction in the modulation of the drift rate by Q-value differences. Taken together, our findings highlight the power of computational analyses using RLDDMs [34]: model-based decomposition of RT distributions revealed substantial group differences in component processes underlying reinforcement learning and action selection, despite the fact that overall RTs were similar between groups.

These results might provide some insights into potential mechanisms underlying the development and maintenance of gambling behavior. In animal models, exposure to uncertainty gives rise to behavioral and neural effects similar to those observed during repeated exposure to drugs of abuse [61–64], conceptually linking behavioral addictions [65] and theories of substance-use-disorders such as incentive sensitization theory [66]. In structured environments, overall experienced uncertainty is inversely related to learning performance, and midbrain dopamine neurons fire maximally during uncertain reward prediction [67]. Likewise, human subcortical dopaminergic structures encode risk [68], and striatal dopamine release in gambling disorder is highest under conditions of maximum uncertainty [69]. There is also some evidence that gambling disorder might be linked to an overall increase in dopamine availability in the striatum [70]. Therefore, one could speculate that an increase in overall uncertainty and concomitant dopamine release [67], combined with a potentially general increase in dopamine levels in the gambling group [70] might underlie the observed effects. In line wth this interpretation, decision threholds / boundary separation parameters were reduced by pharmacologically increasing dopamine levels using the same task reported here [41], and the dopamine precursor tyrosine reduced decision thresholds across two different decision-making tasks [71].

In previous work [23], we examined exploration during reinforcement learning using a restless four-armed bandit task [72] in the same participants. This previous task differs from the present reinforcement learning task in a number of important respects: First, average payoffs of each bandit changed continuously according to gaussian random walk processes, whereas in the present task, reinforcement rates were stable. Second, reward feedback consisted of points in the range of 0-100, whereas in the present task, participants received probabilistic binary (win / no win) feedback. Third, 300 trials in total were performed in the bandit task, whereas the present task was substantially shorter. Finally, in our previous task, a stricter response deadline was included, which precluded us from comprehensively analyzing RTs and RLDDMs. Interestingly, however, in the four-armed bandit task, performance was similar between the gambling and the control group [23]. Yet, computational modeling revealed that they relied less on a “directed exploration” strategy [23] that favours selection of uncertain options for information gain [30]. It is nonetheless striking that in the arguable more complex task in a volatile environment, impairments in the gambling group were arguably more subtle, whereas in the present simpler stationary task, group differences in overall accuracy were substantial. What could account for these relative differences in performance? One possibility is that the required degree of temporal integration plays a role. In the restless bandit task, reward feedback on any given trial provides (almost) complete information on the current value of a chosen bandit (“almost” because outcomes are corrupted by gaussian observation noise). In contrast, in order to accurately estimate the underlying reinforcement rates in the present task, binary outcomes need to be integrated across consecutive trials, which could contribute to the impairments. However, working memory deficits, which might contribute to impairments in feedback integration across trials, are not typical neuropsychological characteristics of gambling disorder [19,73].

FMRI analyses across groups then confirmed 1) a positive correlation between activity in vmPFC and the average Q-value across options, which is in line with a wealth of previous imaging findings [3,4,74,75], including results from the same task [41]. Likewise, reward prediction error effects were replicated in bilateral ventral striatum and ventro-medial prefrontal cortex. However, for both effects, Bayesian analyses revealed at best anecdotal evidence for group differences. Alterations in regions of the reward system, in particular ventral striatum and vmPFC, have frequently been reported in gambling disorder, as outlined in a number of reviews [1,2,76]. However, the directionality of these changes in gambling disorder has long puzzled researchers, as both increases and reductions have been reported [2,5,8–10], e.g. depending on task phases [2,77], contextual factors [5,7] or reinforcer categories [6,78]. Although, numerically, the contrast between positive and negative prediction errors in bilateral striatum appeared somewhat more pronounced in the gambling group, consistent with some earlier observations [2], evidence was only anecdotal (1 < BF_incl_ < 3).

A number of limitations of the present study need to be acknowledged. First, the sample size was relatively small, and findings thus require replication in larger samples. This might be particularly problematic for the fMRI analyses, as trial numbers were relatively low. Second, as is often the case in studies on gambling disorder, due to the greater prevalence of problem gambling behavior in males [79], only male participants were tested, limiting the generalizability of our results. Third, in contrast to the original study according to which the task was set up [43], we only included a gain condition, and no loss condition. The degree to which the reported impairments in reinforcement learning in the gambling group extend to tasks with an explicit loss condition therefore remain to be examined in future studies. Finally, a classification of individuals suffering from disordered gambling into different subtypes according to clinical characteristics, disorder trajectories and/or gambling motivations have been proposed [80,81]. These factors potentially reflect important individual differences in the context of disordered gambling, and the same holds for the preferred gambling format of individuals. However, given the small sample size, examination of such subtypes as well as effects of preferred gambling format is not feasible.

Taken together, here we provide a comprehensive model-based analysis of computational mechanisms underlying impaired reinforcement learning performance in gambling disorder. Model-based decomposition of RTs revealed that, although overall RTs were similar between groups, the underlying processes differed considerably. In particular, the gambling group showed shorter non-decision times, an increasing focus on speed vs. accuracy over the course of the experiment (reduction of boundary separation over time) and a reduced impact of Q-value differences on the drift rate. These findings highlight that reinforcement learning impairments in gambling disorder are likely attributable to alterations in multiple component processes.

## Acknowledgements

This work was funded by Deutsche Forschungsgemeinschaft (PE 1627/5-1 to J.P.). J.P. and A.W. designed the study. A.W. acquired the data. J.P. and A.W. analyzed the data. J.P. wrote the paper. A.W. provided revisions. J.P. supervised the project.

## Data and code availability

Raw data cannot be shared publicly because participants did not provide consent for having raw data posted in a public repository. Raw choice are available from zenodo.org (link will be provided) for researchers meeting the criteria for access to confidential data. Processed fMRI data (T-maps and extracted parameter estimates) for the effects shown in Figures 6 and 7 are available on the Open Science Framework (https://osf.io/mb8zr/). JAGS model code is likewise available on the Open Science Framework (https://osf.io/mb8zr/).

## Supplemental material

**Supplemental Figure 1.**
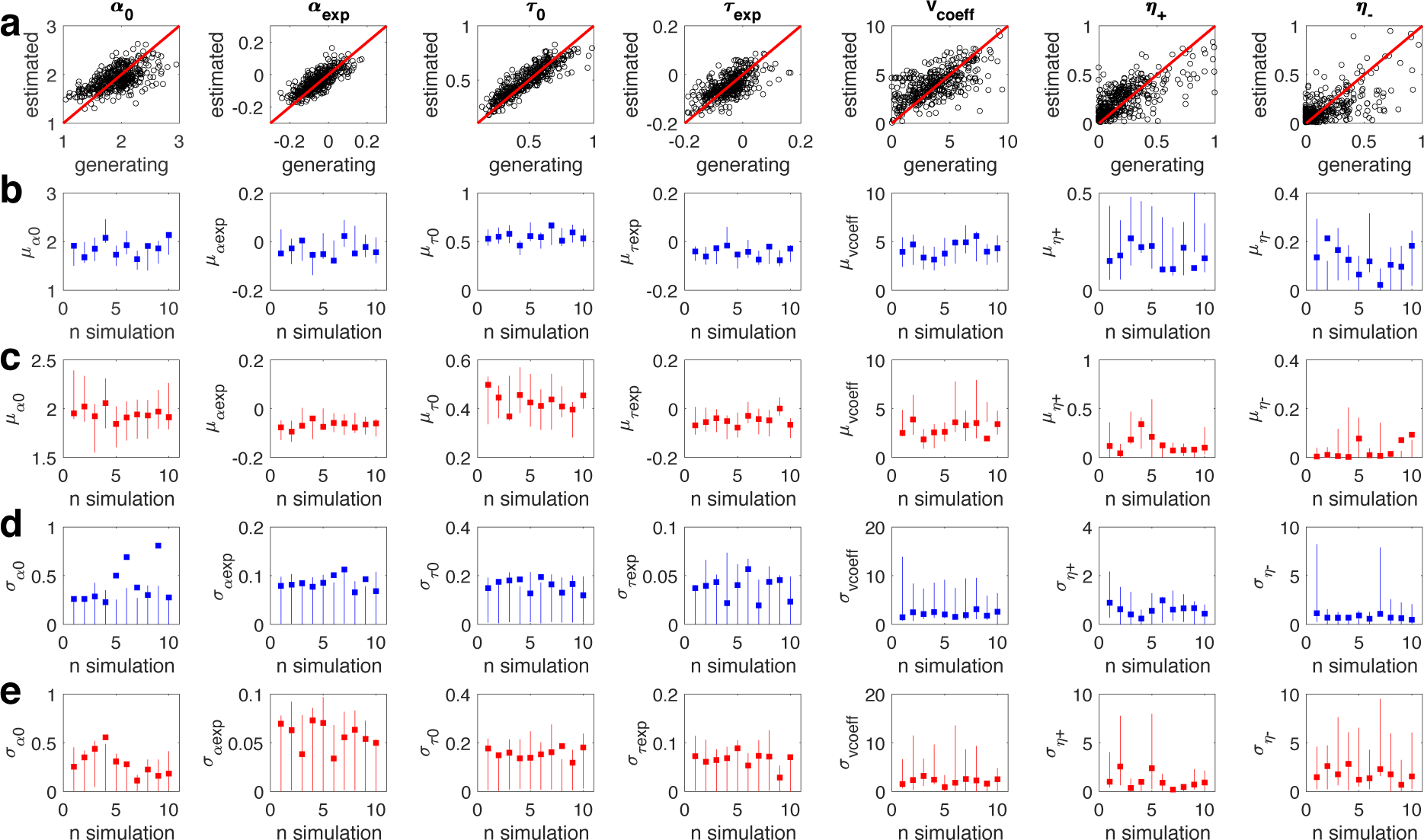
Parameter recovery simulation results for RLDDM8 (dual learning rates, modulated boundary separation and non-decision time). a) Generating vs. estimated single-subject parameters across all 10 simulations. b) Control group parameter means. c) Gambling group parameter means. d) Control group parameter standard deviations. e) Gambling group parameter standard deviations. In b-e, squares denote the generating parameter, and vertical lines denote the 95% highest posterior density of the parameter estimation.

**Supplemental Figure 2.**
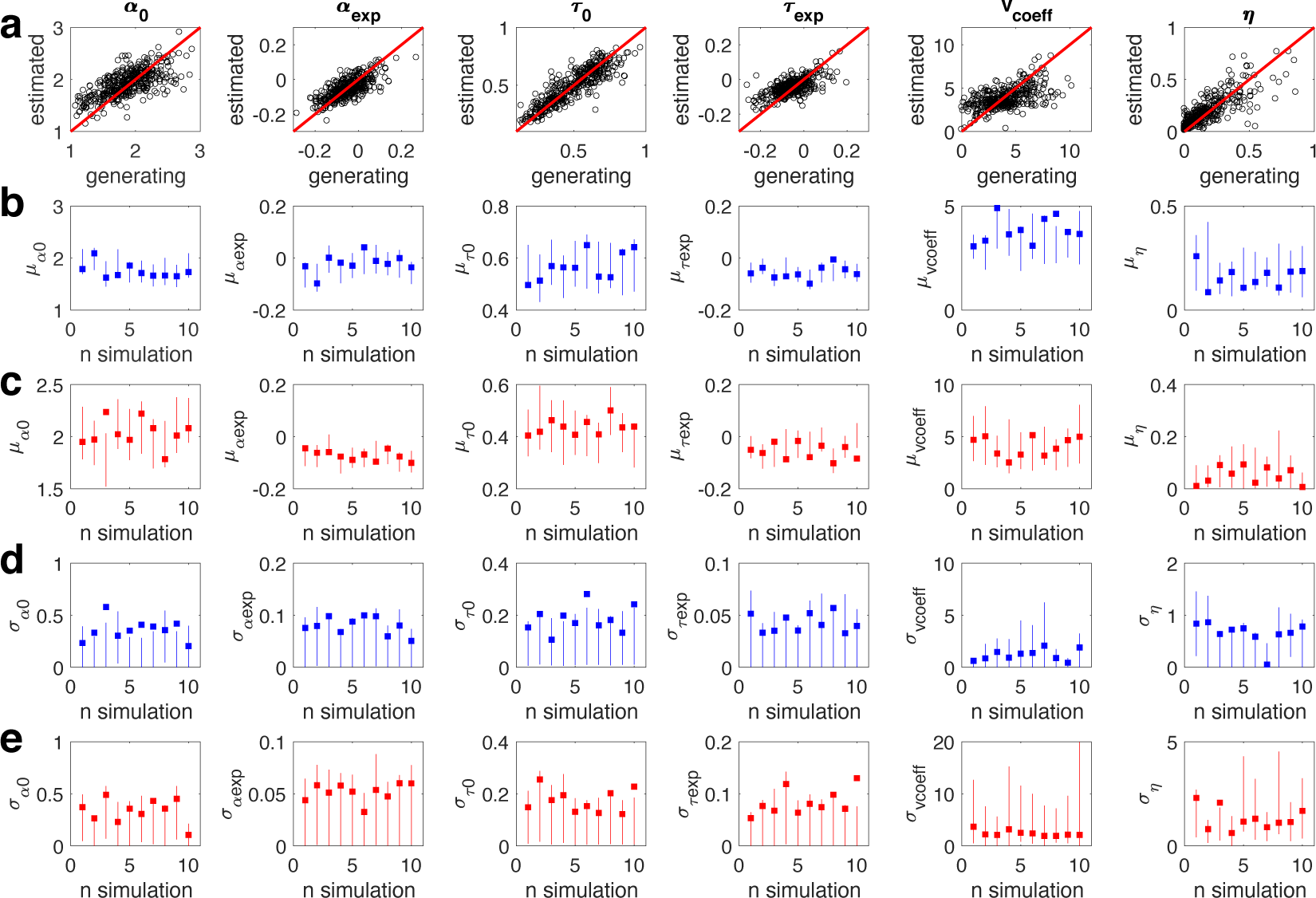
Parameter recovery simulation results for RLDDM4 (single learning rate, modulated boundary separation and modulated non-decision time). a) Generating vs. estimated single-subject parameters across all 10 simulations. Panels b-e show generating group-level parameters (means and standard deviations) plotted as squares, and estimated 95% highest posterior density intervals as vertical lines. b) Control group parameter means. c) Gambling group parameter means. d) Control group parameter standard deviations. e) Gambling group parameter standard deviations.

**Supplemental Table 1.** Pearson correlations between generating and estimated individual-participant parameters (pooled across 10 simulations) for the RLDDM4 and RLDDM8.

| Parameter | RLDDM4 | RLDDM8 |
| --- | --- | --- |
| $\alpha_0$ | .70 | .59 |
| $\alpha_{\text{exp}}$ | .73 | .78 |
| $\tau_0$ | .88 | .90 |
| $\tau_{\text{exp}}$ | .65 | .64 |
| $v_{\text{coeff}}$ | .57 | .69 |
| $\eta$ | .83 | - |
| $\eta_+$ | - | .71 |
| $\eta_-$ | - | .69 |

**Supplemental Figure 3.**
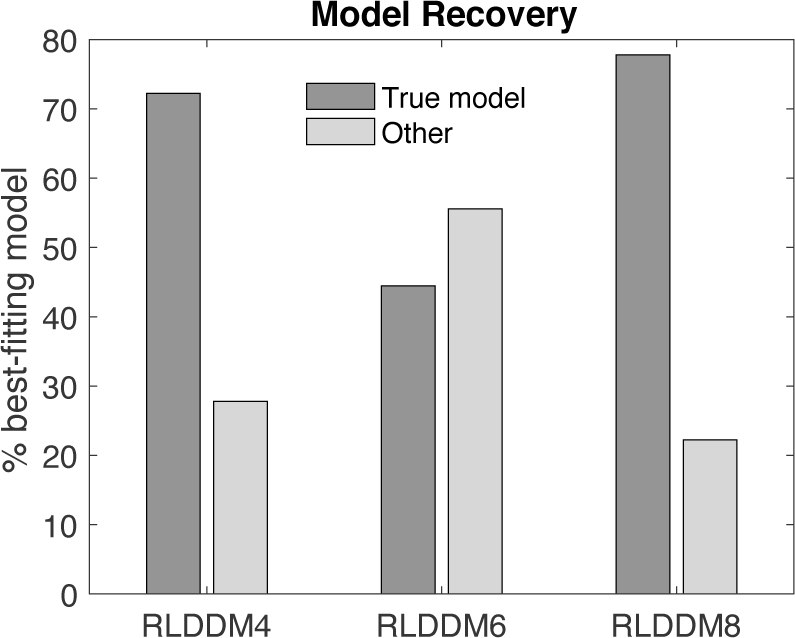
Model recovery results. In addition to the best-fitting model (RLDDM8), model recovery focused on those models that exhibited overlap with RLDDM8 in terms of the 95%CI of the -elpd score in at least one group. This was the case for RLDDMs 4 and 6. We simulated n=20 full datasets from each of the three models, and re-fit the simulated data with all nine models from our model space. Plotted is the percentage of simulations in which the true data-generating model was recovered (*True model*) and the percentage of simulations in which some other model accounted for the data best (*Other*). Recovery was successful in > 70% of simulatons for both RLDDM4 and RLDDM8, whereas it was <50% for RLDDM6. Note that chance level is 11.11%.

**Supplemental Figure 4.**
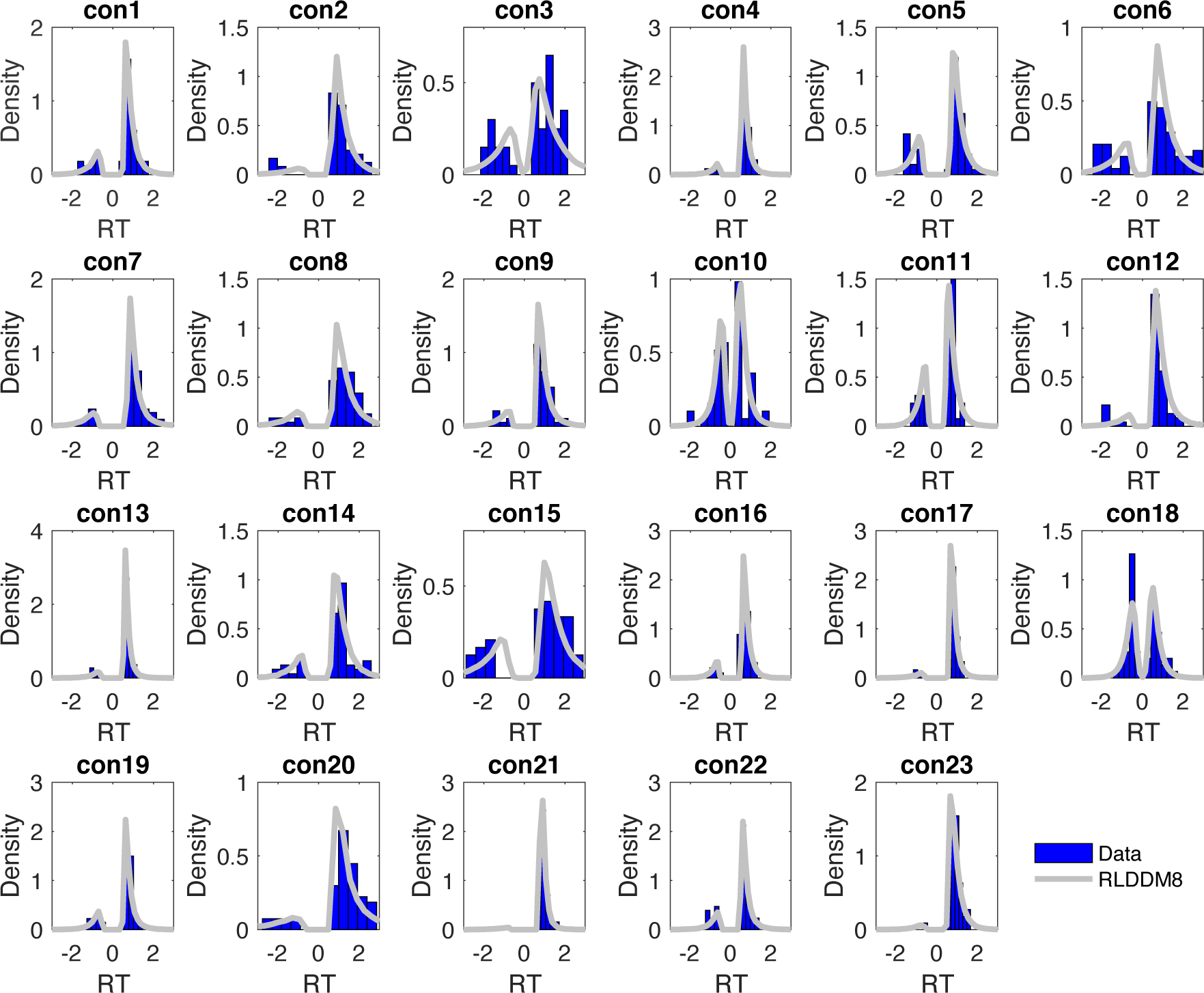
Individual subject posterior predictive checks for the control group and RLDDM8. Plotted are individual-subject observed RT distributions (blue histograms) and model simulated RT distributions (grey lines, smoothed histograms of 1k RT distributions simulated from the model’s posterior distribution.

**Supplemental Figure 5.**
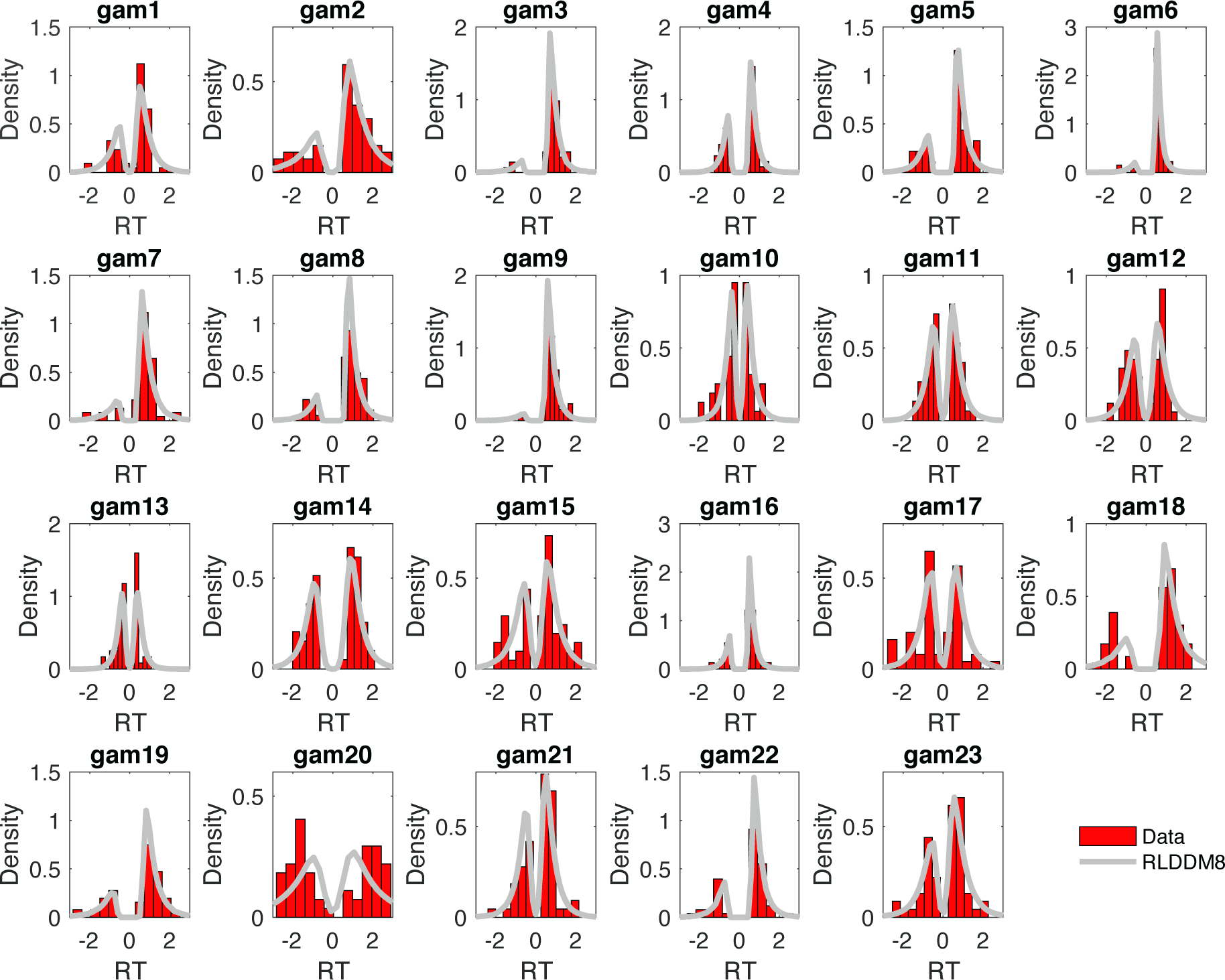
Individual subject posterior predictive checks for the gambling disorder group and RLDDM8. Plotted are individual-subject observed RT distributions (red histograms) and model simulated RT distributions (grey lines, smoothed histograms of 1k RT distributions simulated from the model’s posterior distribution.

**Supplemental Figure 6.**
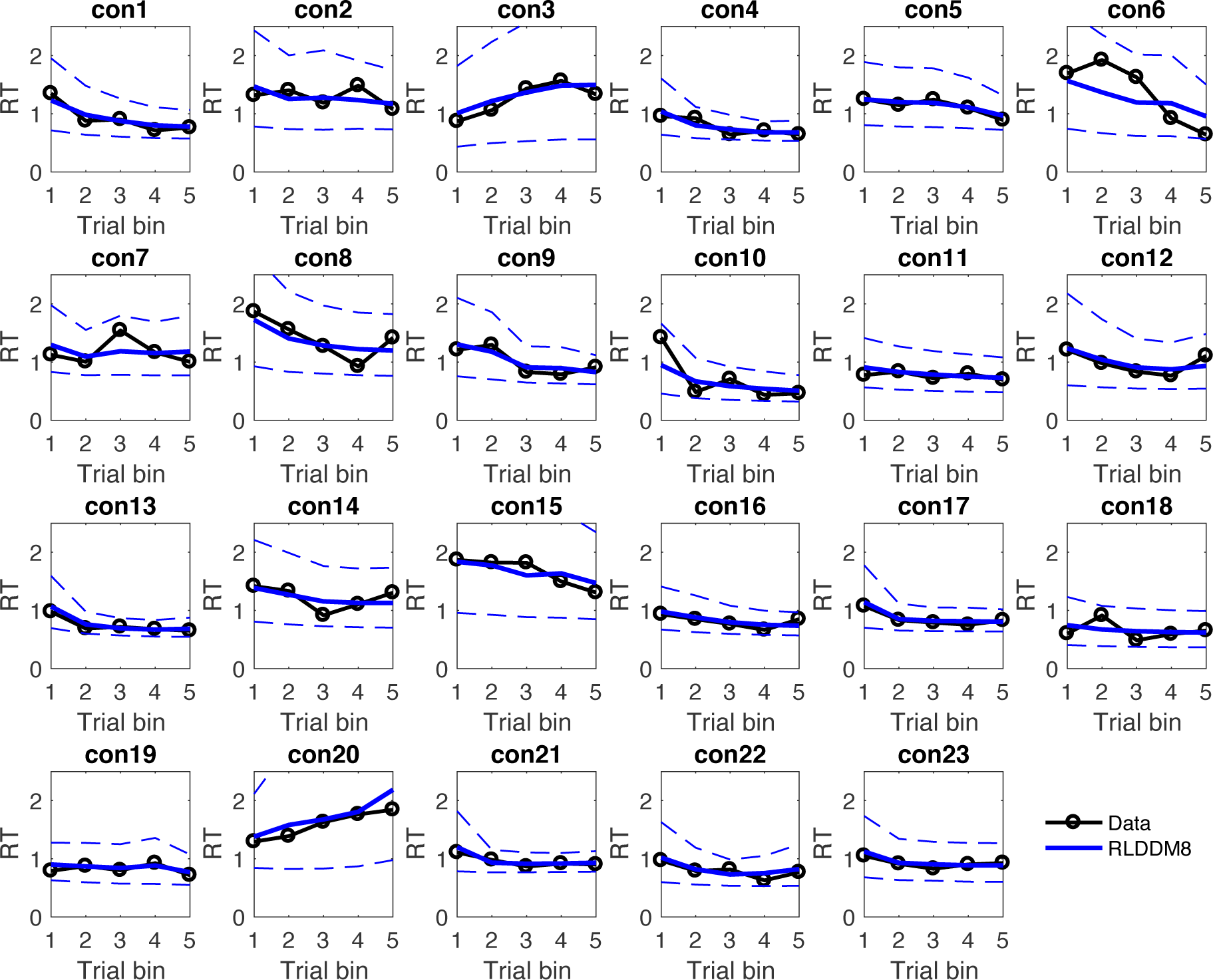
Posterior predictive checks for RT changes over the course of learning in individual control group participants. Black lines denote observed mean RTs per trial bin. Solid blue lines denote mean RTs across 1k simluated data sets from the RLDDM8 posterior distribution. Dashed lines denote the +/-95% percentile of the simulated RTs.

**Supplemental Figure 7.**
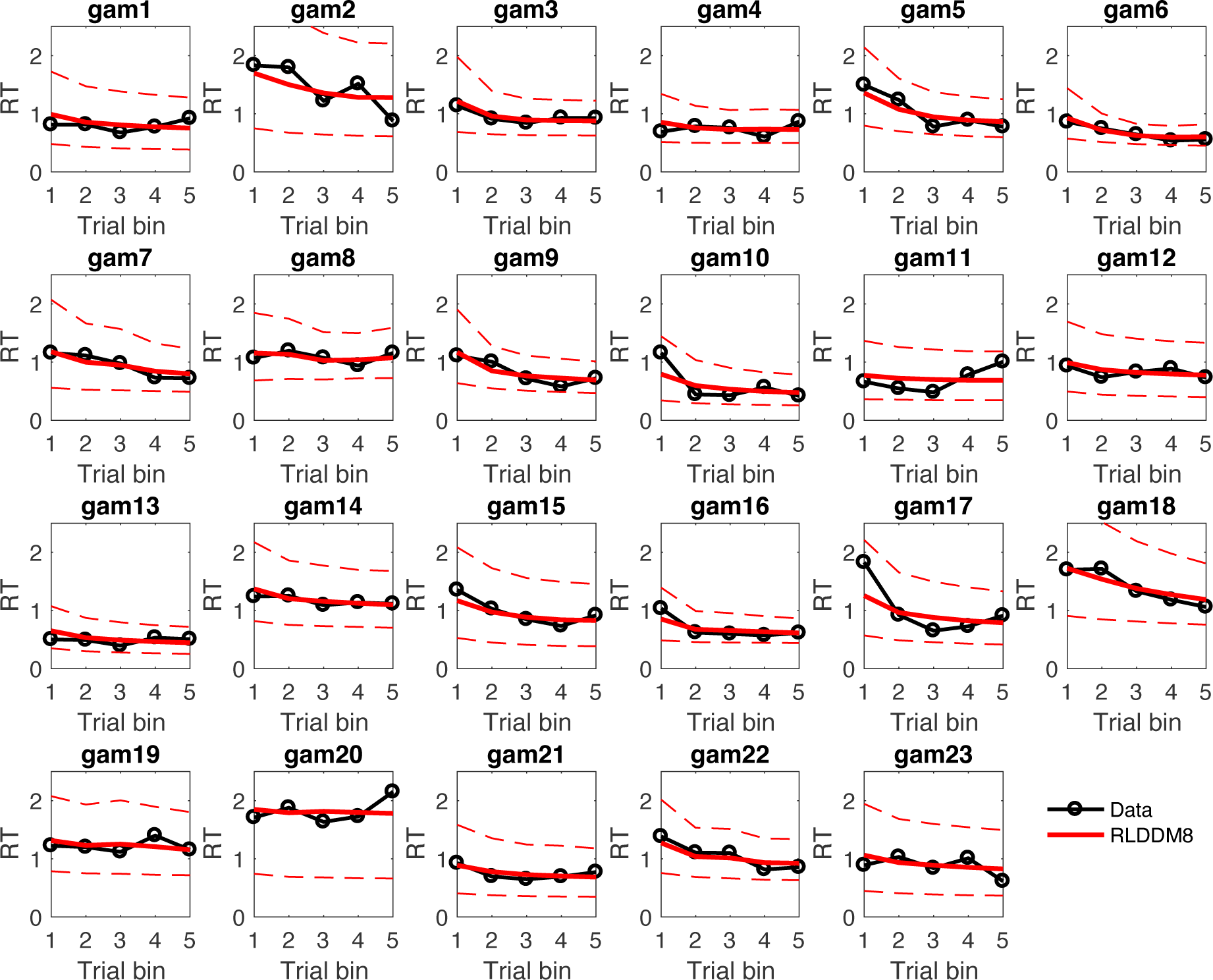
Posterior predictive checks for RT changes over the course of learning in individual gambling disorder group participants. Black lines denote observed mean RTs per trial bin. Solid red lines denote mean RTs across 1k simluated data sets from the RLDDM8 posterior distribution. Dashed lines denote the +/-95% percentile of the simulated RTs.

**Supplemental Figure 8.**
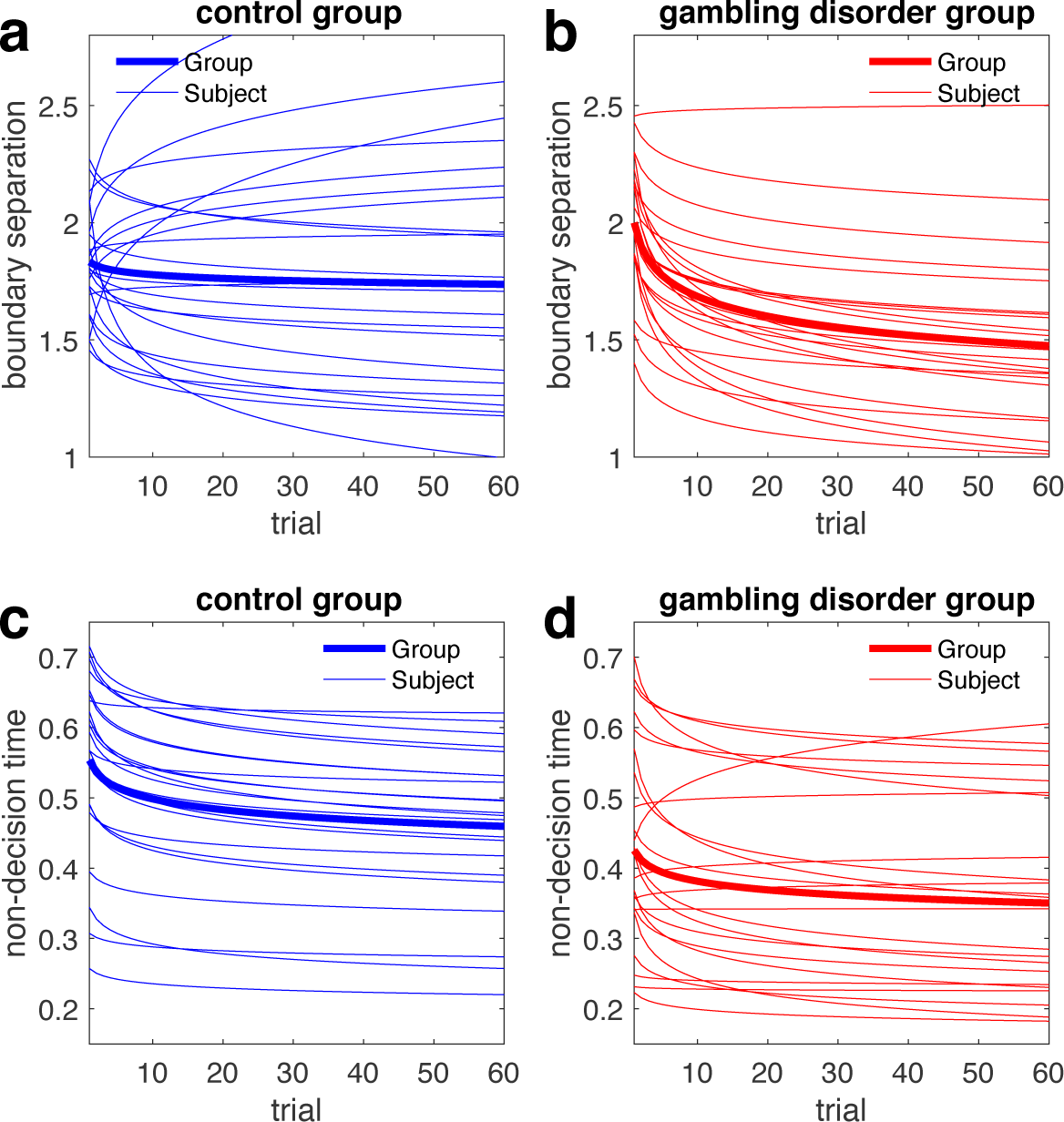
Estimated trial-wise boundary separation (a,b) and non-decision time parameters (c,d) for control group participants (blue) and participants from the gambling group (red) according to RLDDM8. The solid lines plot parameter changes based on the mean group-level posteriors, whereas the thin lines depict individual subject curves.

**Supplemental Figure 9.**
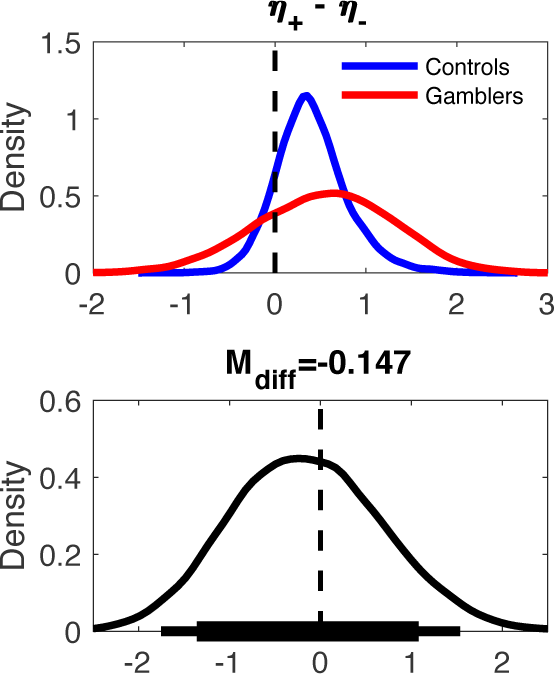
Upper panel: Posterior differences between positive and negative learning rates per group (controls: blue, gamblers. red). Lower panel: posterior group difference in learning rate differences (controls – gamblers). Solid (thin) horizontal lines in the lower panel denotes 85% (95%) highest posterior density interval.

**Supplemental Table 2.**
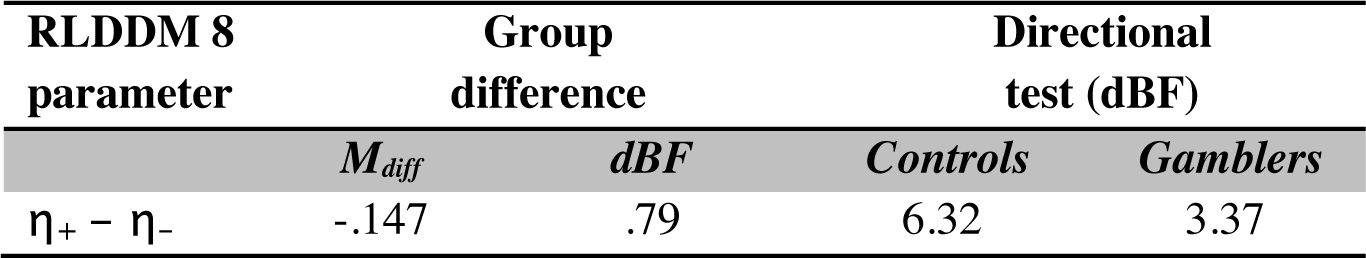
Learning rate differences in the dual learning rate model RLDDM8 (mean posterior group difference (M_diff_) and Bayes factors testing for directional effects, dBF). dBF values > 1 quantify the degree of evidence for a reduction in a parameter in gamblers vs. controls compared to the evidence for an increase. dBF values < 1 reflect the reverse. Directional test refers to tests for directional effects of the learning rate difference performed separately per group.

**Supplemental Figure 10.**
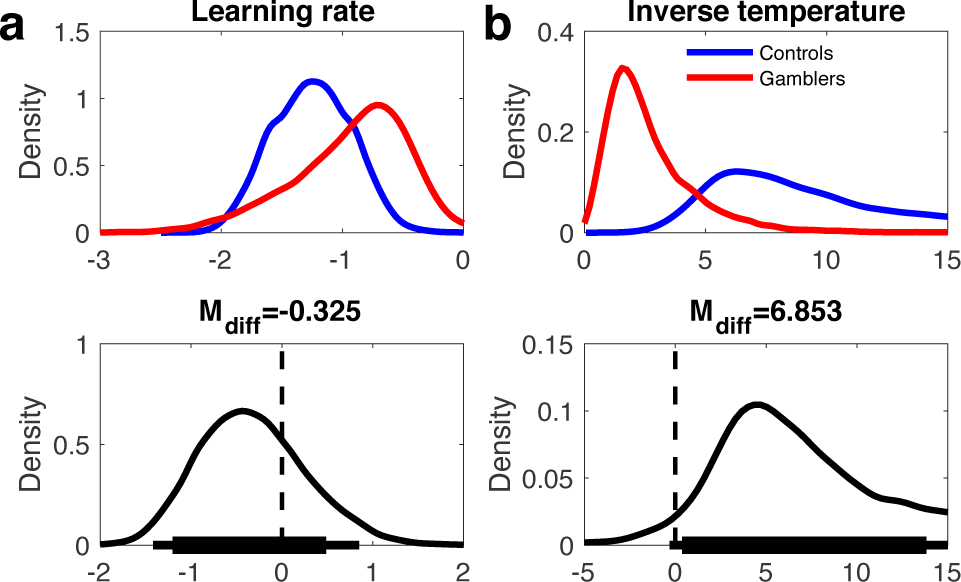
Upper panels: Softmax model posterior distributions of group mean learning rates (a) and softmax inverse temperatures (b) for controls (blue) and gamblers (red). Lower panels: posterior group differences per parameter (controls – gamblers). Solid (thin) horizontal lines in the lower panels denote 85% (95%) highest posterior density intervals. Note that learning rates were fitted in standard normal space [-3, 3] as plotted here, and were back-transformed to the interval [0, 1] via the inverse cumulative normal distribution function.

**Supplemental Table 3.**
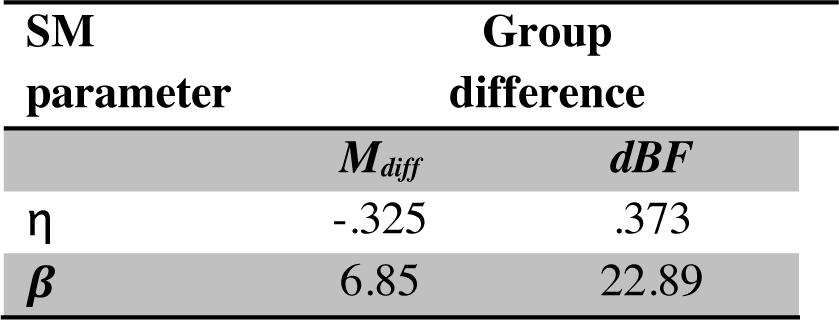
Group differences: Mean posterior group differences in model parameters (M_diff_) and Bayes factors testing for directional effects (dBF). dBF values > 1 quantify the degree of evidence for a reduction in a parameter in gamblers vs. controls compared to the evidence for an increase. dBF values < 1 reflect the reverse.w

| SM<br>parameter | Group<br>difference |  |
| --- | --- | --- |
| | $M_{diff}$ | $dBF$ |
| $\eta$ | -.325 | .373 |
| $\beta$ | 6.85 | 22.89 |

